# Dufour’s gland analysis reveals caste and physiology specific signals in *Bombus impatiens*

**DOI:** 10.1101/2020.06.20.163089

**Authors:** Nathan T. Derstine, Gabriel Villar, Margarita Orlova, Abraham Hefetz, Jocelyn Millar, Etya Amsalem

## Abstract

Reproductive division of labor in insect societies is regulated through multiple concurrent mechanisms, primarily chemical and behavioral. Here, we examined if the Dufour’s gland secretion in the primitively eusocial bumble bee *Bombus impatiens* signals information about caste, social condition, and reproductive status. We chemically analyzed Dufour’s gland contents across castes, age groups, social and reproductive conditions, and examined worker behavioral and antennal responses to gland extracts.

We found that workers and queens each possess caste-specific compounds in their Dufour’s glands. Queens and gynes differed from workers based on the presence of diterpene compounds which were absent in workers, whereas four esters were exclusive to workers. These esters, as well as the total amounts of hydrocarbons in the gland, provided a separation between castes and also between fertile and sterile workers. Olfactometer bioassays demonstrated attraction of workers to Dufour’s gland extracts that did not represent a reproductive conflict, while electroantennogram recordings showed higher overall antennal sensitivity in queenless workers. Our results demonstrate that compounds in the Dufour’s gland act as caste- and physiology-specific signals and are used by workers to discriminate between workers of different social and reproductive status.

## Introduction

Reproductive division of labor in insect societies is regulated through multiple concurrent mechanisms. These often involve a combination of interactions with brood^1,2^, agonistic and policing behaviors among nestmates^3–5^, and chemical signaling among individuals in the colony^6–8^. In the latter, chemical signals from the queen can affect worker physiology and behavior by reducing their ovarian activation^9–12^ and by marking eggs to indicate their maternity, which prevents their consumption by “policing” workers^13–16^. In workers, chemical signals can indicate dominance and fertility status^17–19^, which increases aggressive behavior towards fertile workers^20^ or indicate sterility, which appease aggressive individuals^8,21^. Across social insects, these signals are chemically diverse and have multiple glandular origins^22–24^.

In female Hymenoptera, one such glandular origin is the Dufour’s gland (DG), which is associated with the sting apparatus in females and produces a remarkable diversity of compounds with myriad functions^25,26^. In solitary bees, the DG secretion functions primarily in waterproofing brood cells^27,28^, although there is also evidence that it is the source of a sex pheromone in two parasitoids^29^, and may serve as a nest-marking pheromone in certain solitary bee species^30,31^. In social bees, analyses of the DG secretion have suggested additional functions in social communication^32^, such as signaling of reproductive status and the possible regulation of worker aggressive and foraging behaviors in honey bees and bumble bees^8,21,33,34^.

Bumble bees provide an excellent opportunity to study the role of chemical signals in the regulation of reproduction because they exhibit characteristics of both simple and highly eusocial insect societies. In the first part of the colony cycle, reproduction is dominated by a single queen, with workers being functionally sterile. This changes in the second half of the colony cycle, the competition phase, where workers initiate egg laying and produce males in competition with the queen^35^. In *Bombus terrestris*, the DG secretion varies with caste, reproductive state, age, and task^8,34^. In queens, the DG contains aliphatic saturated and unsaturated hydrocarbons, whereas in workers these are accompanied by long chain esters. The reduction of these esters to non-detectable amounts in queenless workers with activated ovaries suggests that they may signal sterility^8^. There is also a negative correlation between the amount of esters a worker possesses and the aggression directed toward her during worker-worker competition over reproduction, suggesting an appeasement effect of these esters, in line with their hypothesized function as sterility signals^21^. It is unknown if these signals are widespread in *Bombus*, or represents a unique chemical signaling mechanism in *B. terrestris*. The ester amount in *B. terrestris* also increases with age in a similar fashion as the increase in worker ovarian activation and is therefore an important confounding factor to evaluate when examining the glandular compounds and their relation to social and reproductive status^8^. In this study, we leverage our understanding of *B. terrestris* chemical ecology in order to glean new insights into potential shared or elaborated mechanisms mediating social organization across bumble bee species.

To examine the role of the gland from a signal production standpoint and discover if *B. impatiens* females produce a social signal in the DG linked to their fertility status, we examined the chemical composition of the secretion as a function of caste, age, social, and reproductive status. To examine the signaling role of the gland from a sensory standpoint, we tested the behavioral and antennal responses of workers, either queenless or queenright, to the DG secretion of either queenless or queenright workers. The DG may carry multiple social signals which are caste or social status dependent. Depending on what these roles are, DG compounds may induce attraction or avoidance to facilitate cooperation or competition. Therefore, we hypothesize that if workers produce a cooperative signal (e.g., sterile workers producing esters, like in *B. terrestris*), a DG extract of these workers will elicit higher attraction by cooperative workers in comparison with a control extract, and if workers produce a fertility signal, a DG extract of these workers will elicit avoidance in competing workers as compared to control. Additionally, we hypothesize that the secretion from workers carrying a cooperative or competitive signal will elicit stronger antennal responses in the workers receiving these signals, reflecting increased sensitivity towards compounds that may impact their fitness.

## Materials and Methods

### Bee colonies and sample preparation for chemical analysis

Bees were collected from *B. impatiens* colonies obtained from Koppert Biological Systems (Howell, Michigan, USA) or Biobest Canada Ltd. (Leamington, ONT). Upon arrival to the laboratory, colonies were approximately 2-4 weeks old (based on first worker emergence), with less than 30 workers each, a queen, and all stages of brood. Colonies were maintained in nest-boxes at a constant temperature of 28–30°C and 60% relative humidity, constant darkness and supplied *ad libitum* with a 60% sugar solution and honeybee-collected pollen. These colonies were used as sources of newly-emerged workers (<24 h old) that were sampled prior to sexual production phase (i.e., gynes and males). Newly-emerged workers were collected from 7 colonies (see Table S1 for sample size) assigned to one of three treatments: queenright (QR, n=70), queenless (QL, n=70), and queenless and brood-less (QLBL, n=70) workers. QR workers were individually labeled on the day of emergence and remained in their natal colony in the presence of the queen until the day of collection while QL and QLBL workers were housed in a plastic cage (11 cm diameter × 7 cm height) in groups of 3-6 workers without a queen. Queenless groups of workers typically lay eggs within 6-8 days^36^, and the presence of brood may also affect worker reproduction^2^. Therefore, in the QL groups, eggs laid by workers remained in the cages, whereas in the QLBL groups, eggs laid by workers were removed daily. QR workers were collected from young queenright colonies showing no signs of competition over reproduction between the queen and the workers. Workers ranging in age from one to fourteen days old (5/age group) were collected from all treatments. All workers were collected and stored at −80° C until dissection.

Active mated, egg-laying queens (hereafter, “queens”; n=20) were obtained from full-sized colonies with >100 workers. These queens were actively producing female workers prior to sampling. Unmated queens (hereafter “gynes”; n=20) were collected from 3 colonies (see Table S1 for details). They were separated from their natal colonies upon emergence to prevent mating, housed in small cages in groups of 3-5 gynes and sampled at 4 age points: 3, 6, 10, and 14 days after emergence (5/age group).

Olfactometer and electrophysiology bioassays were conducted using 7-day old QR and QLBL workers exposed to DG extracts of either 7-day old QR or QLBL workers, based on differences observed in the chemical analysis and ovarian activation in these groups. Olfactometer bioassays were conducted using 122 workers in total taken from 11 colonies and electrophysiology bioassays were conducted using 29 workers taken from 3 colonies (see Table S1 for breakdown per treatment).

#### Dissection of bees

Ovaries were dissected under a stereomicroscope in distilled water, and the largest three terminal oocytes across both ovaries (at least one from each ovary) were measured with an eyepiece micrometer. The mean of these three oocyte measurements was used as an indicator of ovary activation for each bee. Mean terminal oocytes (hereafter, “oocyte size”) was used as a continuous variable in Figures 2a and 4a. In order to perform the LDA analysis in Fig. 3 we had to categorize bee oocyte size as LDA cannot be performed using continuous variables. We therefore categorized workers into three groups based on their terminal oocyte size: inactive (<0.5 mm), intermediate (0.5 – 2 mm), and active (>2 mm).

**Figure 1.**
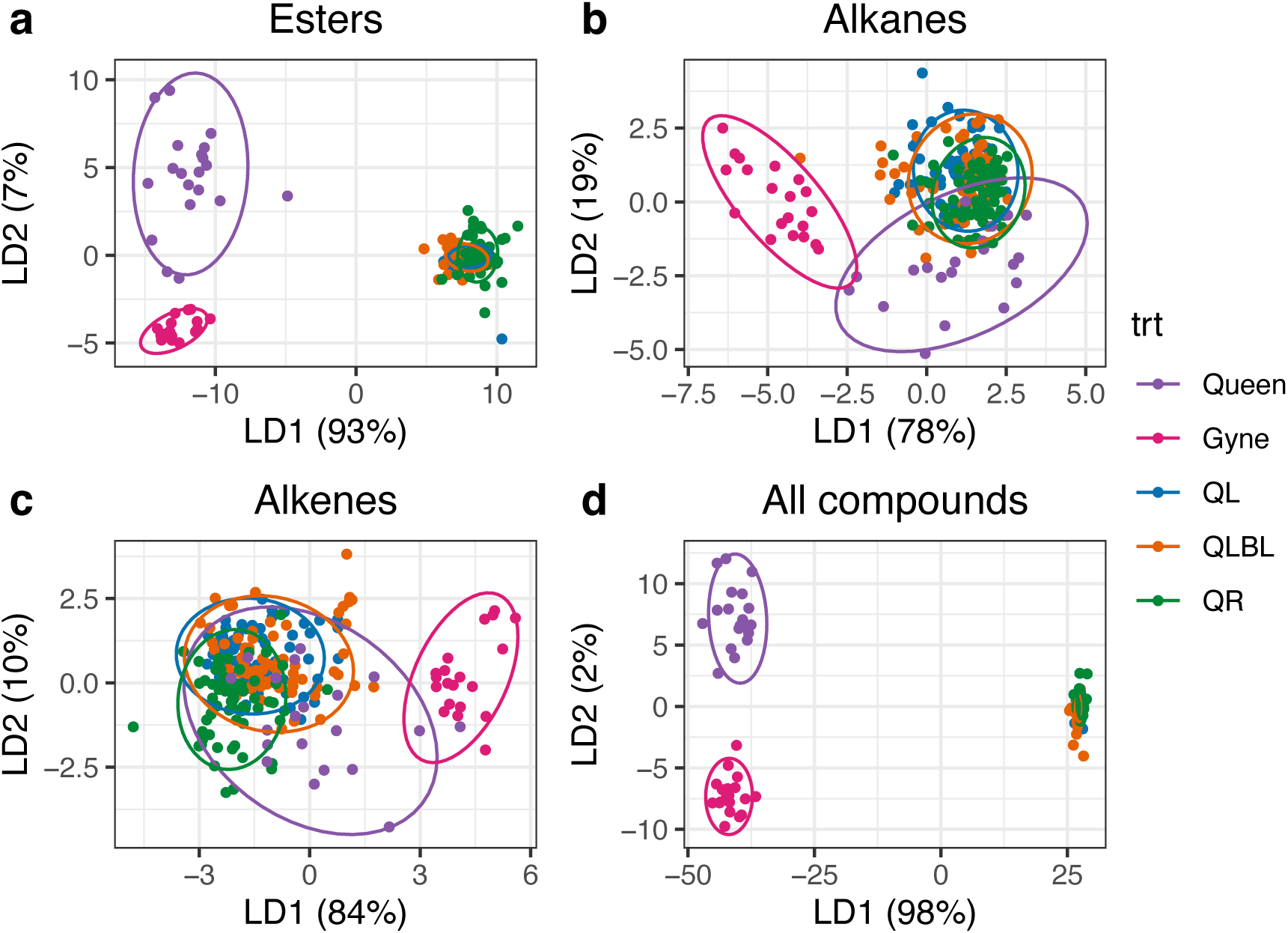
Linear discriminant analyses on Aitchison’s Z-transformed relative peak areas of Dufour’s gland extracts of workers of three social conditions (queenright – QR, queenless - QL, and queenless, brood-less - QLBL), gynes, and queens, with caste/treatment as the grouping variable. Each point is an individual bee. Panels **a-d** show the clustering generated by plotting the first two linear discriminant functions using the esters, alkanes, alkenes, or all compounds.

**Figure 2.**
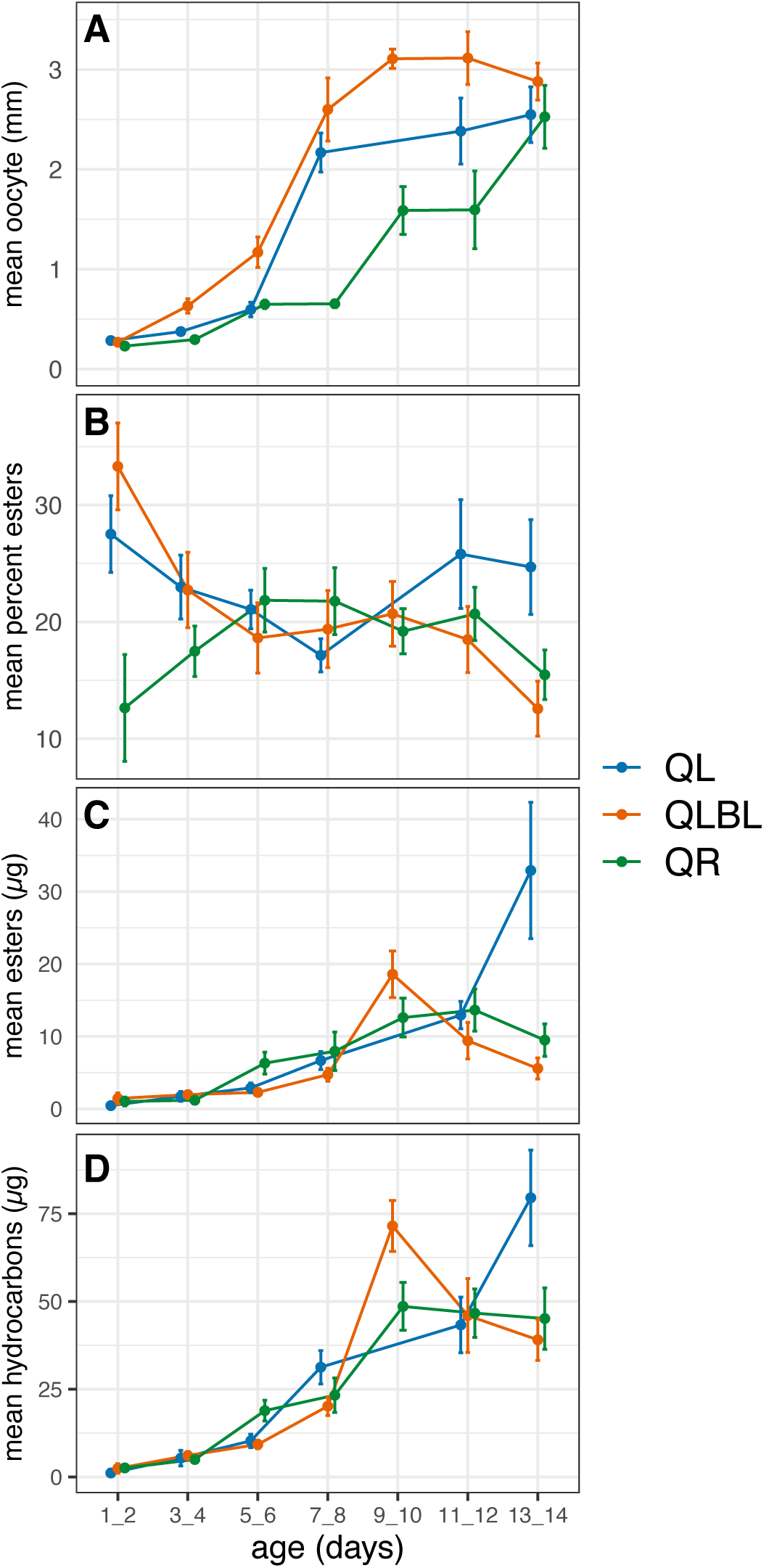
Change in worker reproductive state and Dufour’s gland chemistry as a function of age (1-14 days post emergence) and social condition: queenright (QR), queenless (QL), and queenless, brood-less (QLBL) workers. (A) mean terminal oocyte size (mm), (B) mean percent of total esters, (C) mean µg of total esters, and (D) mean µg of total hydrocarbons. Data are grouped into two-day bins and represent the mean value of 10 workers per age group ± S.E.M.

**Figure 3.**
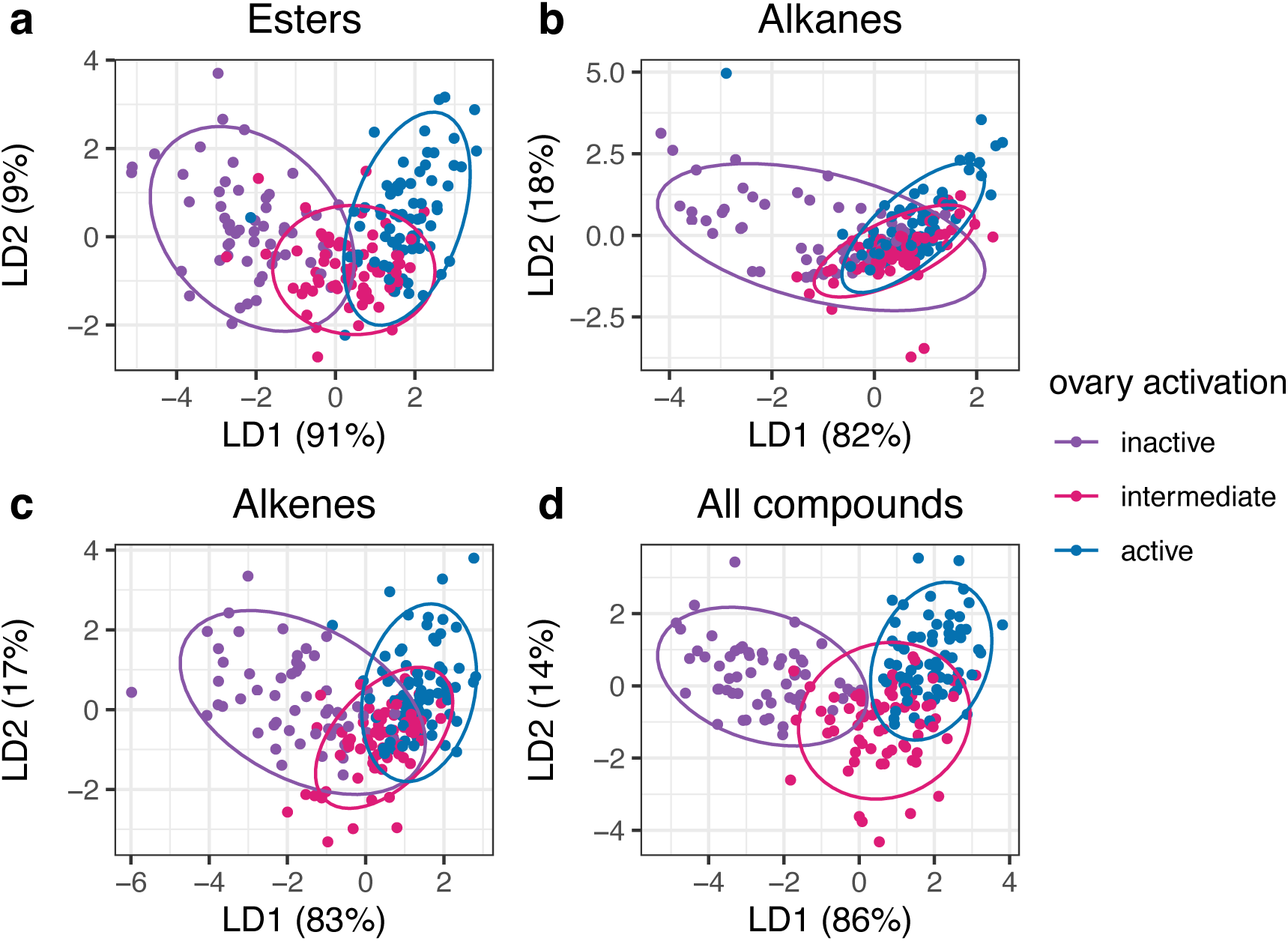
Linear discriminant analyses on Z-transformed relative peak areas of Dufour’s gland extracts of workers with three ovarian activation categories as the grouping variable. Each point is the value from an individual bee. Panels **a-d** show the clustering generated by plotting the first two linear discriminant functions using the esters, alkanes, alkenes, or all compounds.

**Figure 4.**
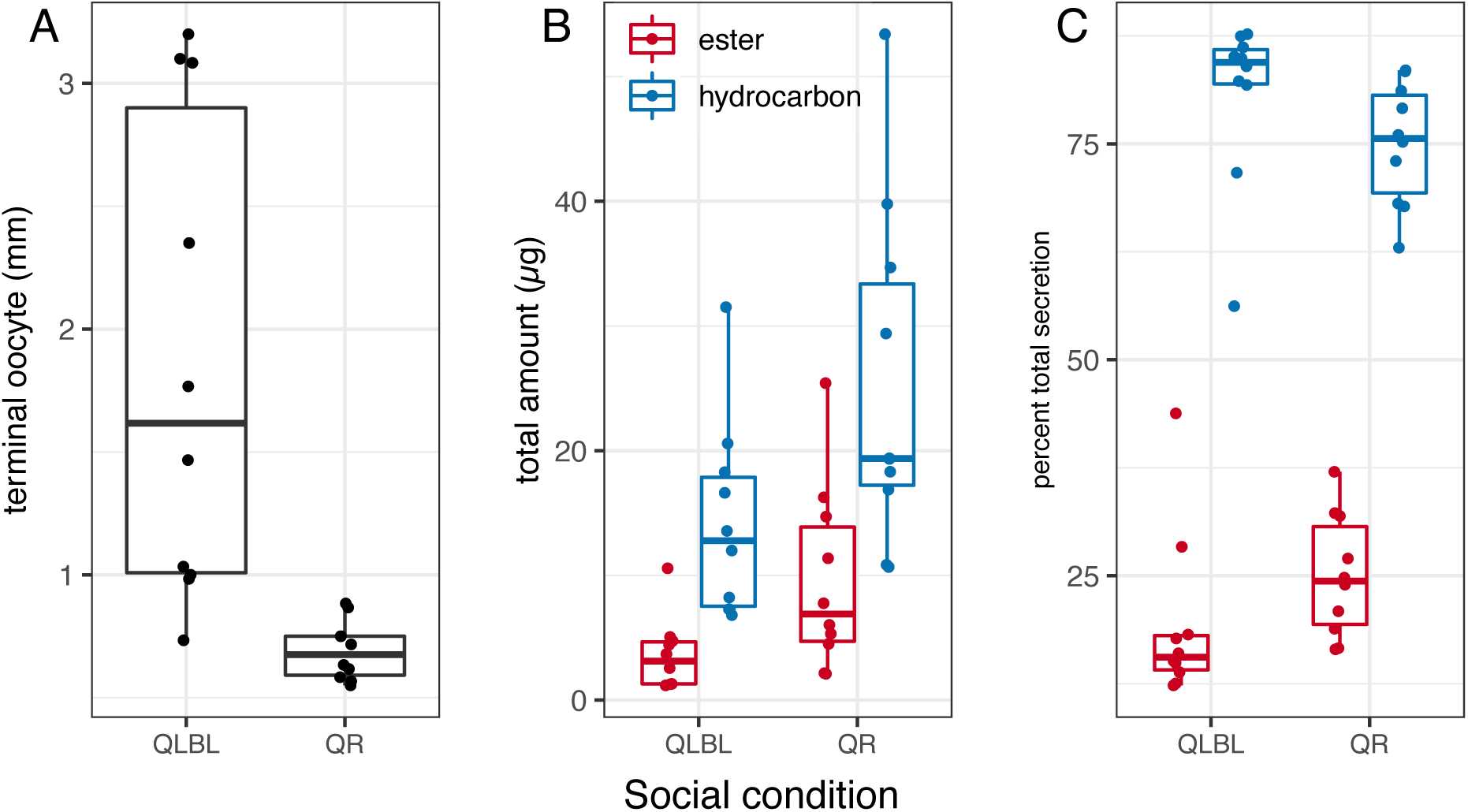
Reproductive state and Dufour’s gland chemistry of 6-7 day old workers in queenless (QR) and queenless-broodless (QLBL)groups. (A) mean terminal oocyte size (mm), (B) mean µg of total esters and hydrocarbons ± SE (n = 10), and (C) mean percent of total esters and hydrocarbons ± SE (n = 10). Each data point is the value from an individual worker.

DGs from each individual were dissected and immediately immersed in hexane. Glands were extracted for 48 hours after which each sample was evaporated and reconstituted in 100 µL hexane spiked with an internal standard of 1 µg eicosane (C_20_).

#### Chemical analysis

To identify compounds in the DG, extracts were analyzed on two separate mass spectrometers. At UC Riverside, samples were analyzed on a Hewlett-Packard (H-P, now Agilent Technologies, Santa Clara CA, USA) 6890 gas chromatograph (GC) equipped with a DB-17 mid-polarity column (0.25 mm id x 30 m x 0.25 µm film thickness; J&W Scientific, Folsom CA, USA) interfaced to an H-P 5973 mass spectrometer. The temperature program was as follows: 100°C (0 min. hold) to 160°C at 20°C/min, then 4°C/min to 280°C (hold for 25 min). Injections were made in split-less mode, with injector and transfer line temperatures of 280°C. At Penn State, samples were analyzed on an Agilent 7890A GC equipped with a HP-5ms column (0.25 mm id x 30 m x 0.25 µm film thickness, Agilent) and interfaced to an Agilent 5975C mass spectrometer. Compounds were tentatively identified based on diagnostic ions in the resulting spectra and retention indices, and where possible, identifications were confirmed by matching retention times and mass spectra with those of authentic standards.

Compounds in DG extracts were quantified on a Trace 1310 GC (Thermo Fisher, Waltham, MA USA) equipped with a flame-ionization detector (FID) and a TG-5MS column (0.25 mm id x 30 m x 0.25 µm film thickness; Thermo Fisher). The temperature program was as follows: 60°C to 120°C at 15°C/min followed by 120°C to 300°C (5m hold) at 4°C/min. The injector port and FID were held constant at 250°C and 320°C, respectively.

#### Olfactometer Bioassays

To examine if *B. impatiens* workers perceived the DG secretion by olfaction, and if their response depended on reproductive or social state, bioassays were conducted using two-choice olfactometers as previously described^38^. Olfactometers (see Fig. S1 for illustration) were fashioned from plastic petri dishes (150 × 15 mm) with two equidistant holes (2 cm diameter) in the bottom providing access to small plastic cups that held treatment or control stimuli (QR or QLBL DG extract or control). One gland equivalent of extract (100 µl total volume) was pipetted onto glass coverslips and the solvent was allowed to evaporate before being introduced into the cups, which were subsequently affixed to the olfactometer. After lifting the petri dish lid enough to allow entry, workers were placed in the center of the olfactometer and given 60 min to make a first choice, which was recorded. Bees did not have the ability to touch the glass slides unless they fully entered the cup, and were prevented from returning to the main arena after a choice was made via a 3 cm section of plastic straw glued to each hole. Bioassays were conducted under a red light and olfactometers were washed with soap and water and allowed to air dry before being reused. Cups containing test stimuli, extracts and bees were only used once. All behavioral assays were conducted in the rearing chamber between noon to 4 PM. Seven-day old QR or QLBL workers were tested for their preference between 2 choices in three bioassays: QR DG extract vs. callow DG extract, QLBL DG extract vs. callow DG extract, and QR DG extract vs. QLBL DG extract. Overall, we conducted 122 bioassays (16-25 replicates per treatment group and social condition) which resulted in 71.3% response rate. Bioassays where bees did not respond within the time frame were not included in the analysis. Non-responder bees were not associated with any specific treatment (proportion test, χ^2^ = 0.16, p = 0.69, 25.9% and 30.9% of QLBL and QR non-responders, respectively). Callow DG extract was used as control as it is undifferentiated into a specific chemical profile, being mostly devoid of potential signal molecules, but went through the same extraction process as 7-day old worker DG. This captured methodological noise that a solvent control would not. In addition, the preference to a solvent control and callow extract were not significantly different when compared directly (binomial exact test, p = 0.77, n = 12, 20% non-responders, data not shown). In all bioassays, extracts were presented to workers as a dose of 1-bee equivalent. Extracts were prepared as described above for the chemical analyses, pooled, and the concentration was adjusted to 1 gland equivalent per 100 µL hexane.

#### Electroantennogram recordings

Odor cartridges were created using 7-day old worker DG extracts, either QR or QLBL, as follows. 1 DG equivalent per 100 µL of hexane (DG extracts were prepared as above) was administered onto strips of Whatman filter paper. After the solvent had evaporated, the filter paper strips were inserted into glass pipettes and sealed with aluminum foil until use. A “blank” control consisting of 100 µL of solvent was allowed to evaporate from the filter paper before being placed and sealed in the cartridge. For the electroantennograms (QR n=13, QLBL n=16), the right antenna was excised and the tip snipped with a fine scissors prior to mounting on a quadroprobe electroantennogram system with Spectra 360 electroconductive gel^39^. A constant airflow of charcoal-purified humidified air was passed across the antenna through a 10 mm glass tube for the duration of each experiment. Each antenna was allowed to reach a steady baseline before starting the experiment, at which point the test odorants were delivered into a steady airstream using the odor cartridges in conjunction with a stimulus flow controller (Syntech, Hilversum, Netherlands). A 40 ml/s air pulse was pushed through the cartridge for 0.05 seconds, effectively delivering 2 ml of volatiles into the air stream and across the antenna. Responses to each stimulus were amplified using a quadroprobe, 10x pre-amplifier (Syntech, Netherlands) and further amplified 10x before being recorded and analyzed on a desktop computer using custom software (as in^40^). Odorants from control, QLBL, or QR extracts were presented to each antenna 30 seconds apart, after the antennal response had returned to baseline activity. Responses to a final blank puff were subtracted from the responses to the test odorants and the corrected values were used in analyses. Each worker antenna was exposed to control, QLBL, and QR extracts, and each odor cartridge was used twice.

#### Statistical analysis

Statistical analyses were performed using R (version 3.6.1) in RStudio (version 1.2.1335) and SPSS v.21 (IBM) software. To examine differences in the DG secretion between females of differing caste (queen, workers) and social conditions (QR, QL, QLBL worker groups), linear discriminant analyses (LDA) were performed using Aitchinson Z-transformed relative peak areas as predictor variables, and social condition, caste, or age as grouping variables. Only peaks > 0.5% of the total peak area were used. These selected peaks were then re-standardized to 100% and transformed using Aitchinson’s formula [Z_ij_ = ln[Y_ij_/g(Y_j_)]^41^, to reduce collinearity prior to multivariate analysis^42,43^. For peaks that were not present in all groups, a small nominal value was added as the transformation cannot be calculated with zero values. In some cases, poorly separated clusters of peaks prevented accurate integration, and after mass spectral confirmation that these were of the same compound class (e.g. esters), they were integrated together as a “complex” and used as a single variable.

The effects of colony, age, treatment and interactions between them on oocyte size and the amounts of DG compounds were analyzed using SPSS. Neither oocyte size nor absolute amounts of esters, hydrocarbons and total secretion in DG were normally distributed. For this reason, all these parameters underwent a Box Cox transformation prior to being included in the model. Oocyte size was analyzed using Generalized Linear Mixed Models (GLMM) analysis with treatment, age in 2-day intervals and the interaction between them as fixed factors and colony as a random factor. Robust estimation was used to handle violations of model assumptions and Satterthwaite correction was employed to account for small and unequal sample sizes. Absolute amounts of esters, hydrocarbons and total secretion in DG were analyzed using GLMM with treatment, age in 2-day intervals and the interaction between them as fixed factors, oocyte size and its interaction with treatment as covariates, and colony as a random factor. Robust estimation was used to handle violations of model assumptions and Satterthwaite correction was employed to account for small and unequal sample sizes. For all categorical fixed factors a post-hoc contrast analysis with Least Significant Difference (LSD) correction for multiple comparisons was performed. Wilcoxon tests were used to compare oocyte size, ester and hydrocarbon amounts, and ester proportions between QLBL and QR 6 and 7-day old workers. Choice data from olfactometer bioassays were analyzed using exact binomial tests, calculating two-tailed probabilities for a difference in the proportion of choices against the null hypothesis of 50%, or no preference. The average strength of antennal responses from electroantennogram recordings were compared using t-tests when comparing responses based on antennal social condition, and paired t-tests when comparing antennal responses within social condition and between extract type. Significance was evaluated at the α = 0.05 level.

## Results

### Chemistry of *Bombus impatiens* Dufour’s gland

DG extracts and the average oocyte size were analyzed in 210 workers (5/age group, ages 1-14 days for 3 treatment groups), 20 gynes (5/age group, ages 3, 6, 10, 14 days), and 20 queens (4-15 weeks from the emergence of the first worker). During analysis, we discovered 11 outlier worker samples, which were excluded from further analyses. One QR worker had fully activated ovaries at 8 days of age, which was a clear outlier, and 10 QL workers on days 9 and 10 sharing the same 2 cages and from the same colony had unusually undeveloped ovaries which likely resulted from a shared effect related to rearing conditions (e.g., underfeeding) or maternal colony. Thus, 199 workers were included in subsequent analyses. Overall, GC-FID analysis of *B. impatiens* DG extracts in females revealed 98 compounds comprised of aliphatic alkanes (C_21_-C_31_), alkenes, wax esters, and diterpenes (Fig. S2, Table S2). Of these, we included in the statistical analysis 28 compounds that were present in the majority of samples and each constitutes at least 0.5% of the total secretion.

### Examining caste specificity in Dufour’s gland secretion

The DGs of *B. impatiens* females contain compounds that are both queen-specific and worker-specific, in addition to linear alkanes and alkenes found in both castes (Fig S2). Gynes and queens differed from workers based on the presence of the diterpenes β-springene and isomers, which were absent in workers, and long chain esters that were specific to gynes. Conversely, three dodecyl esters (dodecyl octanoate, dodecyl decanoate, and dodecyl dodecanoate), and hexadecyl decanoate were specific to workers (Fig S2, Table S2). In addition, workers produced large quantities of a fifth ester, dodecyl oleate, which was present only in trace amounts in gynes. In gynes and queens, β-springene accounted for 3.2% ± 0.19 and 1.3% ± 0.33 (mean ± SE, n=20), on average, of the total secretion, respectively. The worker-specific esters comprised, on average, 1.0% ± 0.07, 2.6% ± 0.08, 1.8% ± 0.08, 1.5% ± 0.13, and 5.1% ± 0.28 (mean ± SE, n=199) of the total secretion across workers of all ages and treatments, respectively. Long chain esters found in gynes (tentatively identified as terpene esters 6 and 7) comprised, on average, 1.2% ± 0.18 and 2.2% ± 0.26 (mean ± SE, n=20) of the gyne total secretion, while only being detected in trace amounts in workers, and at <1% in DG extracts of queens.

Discriminant analyses using either all compounds or specific chemical classes (esters, alkanes, and alkenes) as predictor variables showed that esters enabled differentiation between queens, gynes, and workers with similar explanatory power as the analyses using all compounds (Fig 1). Figure 1 shows plots of the data according to the first two linear discriminant functions based on the compound class used as predictor variables. Of the worker esters, dodecyl decanoate provided the greatest separation between queen and worker castes along LD1 (coefficient of LD1 1.71, % variation explained = 93%), while of the gyne esters, terpene esters 6 and 7 accounted for the most variability along LD2 (coefficients of LD2 0.94 and −1.22 respectively, % variation explained = 7%).

### Examining the effect of age, social condition, and reproductive state on worker Dufour’s gland secretion

The discriminant analysis data indicate clear separation between queens, gynes, and workers, but produced little to no separation between the worker groups (Fig. 1). However, the large differences between the castes may have masked the more subtle differences between workers. Therefore, we analyzed the three worker groups separately from gynes and queens and tested the effects of worker age, social conditions, and reproductive status on their DG composition. In Fig. 2 we show data pertaining to the overall changes observed in oocyte size, DG esters (absolute and relative amounts) and hydrocarbon absolute amounts in workers as function of their age and social conditions. In the description below we also provide information on the interactions between these variables, and between further variables not shown in Fig. 2 such as the DG total secretion (which is the sum of esters and hydrocarbons). We show the split of hydrocarbons to alkanes and alkenes in Fig. S3.

#### Oocyte size

Oocyte size was significantly affected by social condition and age with a significant interaction between the two (GLMM, F_6,175_ = 73.13, p < 0.001 for age, F_2,179_ = 37.26, p < 0. 001 for social condition, F_11,175_ = 3.44, p < 0.0001 for interaction, Fig. 2A). QLBL workers had larger oocytes than both QL and QR workers (post-hoc LSD pairwise contrast, p < 0.001) and QL workers, on average, had slightly larger oocytes than QR workers (post-hoc LSD pairwise contrast, p = 0.05). Oocyte size started increasing on days 3-4 and reached a plateau on days 7-8 (post-hoc LSD pairwise contrast, p < 0.01 for days 1-2 vs. 3-4, 3-4 vs. 5-6, 5-6 vs. 7-8, p > 0.05 for days 7-8 vs. later time points). Post-hoc analysis of the interaction between social condition and age revealed that oocyte size increased with age similarly in QL and QLBL groups (post-hoc LSD pairwise contrast, p < 0.01 for days 1-2 vs. 3-4, 3-4 vs. 5-6, 5-6 vs. 7-8, p > 0.05 for days 7-8 vs. later time points), while in QR groups the increase started at a later age (5-6 days) (post-hoc LSD pairwise contrast, p > 0.05 for days 1-2 vs. days 3-4 and days 9-10 vs. later time points, p < 0.01 for other comparisons). QLBL workers developed larger oocytes than those in other groups by days 3-4 and the difference persisted at later time points (post-hoc LSD pairwise contrast, p > 0.05 at days 1-2, p < 0.01 for all other comparisons). Based on these differences, we disentangle the effects of age and oocyte size on DG chemistry by including oocyte size as a covariate in all subsequent models.

#### Ester proportion

The relative proportion of DG esters was significantly affected by social condition and age with significant interaction between the two factors (GLMM, F_6,173_ = 3.08, p = 0.007 for age, F_2,172_ = 6.46, p = 0.002 for social condition, F_11,173_ = 2.41, p = 0.008 for age*social condition interaction). Ester proportion covaried with oocyte size (F_1,173_ = 13.06, p < 0.001) with significant interaction between oocyte size and social condition(F_2,175_ = 7.37, p = 0.001) (Fig.2B). Overall, QR workers had lower proportion of esters than QL and QLBL workers (post-hoc LSD pairwise contrast, p = 0.001 for QLBL vs. QR, p = 0.019 for QL vs. QR and p = 0.205 for QL vs. QLBL), but the differences were mostly driven by the early and late timepoints, and the proportions fluctuate significantly over the examined age groups (post-hoc LSD pairwise contrast, p < 0.05 for comparisons of days 1-2 to 7-8 and 9-10 vs. 13-14, p > 0.05 for all other comparisons).

#### Ester amount

The amount of esters in the DG was significantly affected by age but not by social condition, without significant interaction between the two factors, and without covariance with oocyte size or interaction between oocyte size and social condition (GLMM, F_6,173_ = 5.75, p < 0.001 for age, F_2,174_ = 1.41, p = 0.25 for social condition, F_11,172_ = 1.66, p = 0.087 for age*social condition interaction, F_1,175_ = 0.188, p = 0.67 for oocyte size, F_2,174_ = 0.01, p = 0.99 for oocyte size*social condition interaction, Fig. 2C). Post hoc analysis revealed that esters amount increased with age up to days 9-10 (post-hoc LSD pairwise contrast, p < 0.05 for comparisons of days 1-2 to 7-8, p > 0.05 for all later time points).

#### Hydrocarbon amount

The amount of DG hydrocarbons was significantly affected by social condition and age with significant interaction between the two factors. Hydrocarbons amount covaried significantly with oocyte size with significant interaction between oocyte size and social condition (GLMM, F_6,173_ = 9.06, p < 0.001 for age, F_2,174_ = 7.46, p = 0.001 for social condition, F_11,173_ = 2.45, p =0.007 for age*social condition interaction, F_1,175_ = 19.23, p < 0.001 for oocyte size, F_2,174_ = 4.89, p = 0.009 for oocyte size*social condition interaction, Fig. 2D). Overall, QR workers had higher amounts of hydrocarbons than QL and QLBL workers (post-hoc LSD pairwise contrast, p < 0.001 for QLBL vs. QR, p = 0.011 for QL vs. QR and p = 0.111 for QL vs. QLBL). Post hoc analysis revealed that hydrocarbon amounts also increased with age until day 7 at which point a plateau was reached (post-hoc LSD pairwise contrast, p < 0.05 for comparisons of days 1-2 to 7-8, p > 0.05 for all later time points). The analysis of interaction between age and social condition showed that differences between social conditions began on day 5 with the highest amounts of hydrocarbons in QR workers and the lowest amounts in QLBL workers (post-hoc LSD pairwise contrast, p < 0.05 for QR vs. QLBL, p > 0.05 for other comparisons, on days 11-12 p < 0.05 for all comparisons).

#### Total secretion

The amount of DG total secretion was significantly affected by social condition and age with significant interaction between the two factors, and covaried significantly with oocyte size without significant interaction between oocyte size and social condition (GLMM, F_6,173_=8.48, p < 0.001 for age, F_2,173_=4.78, p = 0.01 for social condition, F_11,172_ = 2.06, p = 0.026 for age*social condition interaction, F_1,174_= 12.06, p = 0.001 for covariance with oocyte size, F_2,174_ = 2.53, p =0.082 for oocyte size*social condition interaction, data not shown). Overall, QR workers had higher amounts of total secretion than QL and QLBL workers (post-hoc LSD pairwise contrast, p = 0.002 for QLBL vs. QR, p = 0.032 for QL vs. QR and p = 0.237 for QL vs. QLBL). Post hoc analysis revealed that the total secretion increased with age until day 7 at which point a plateau was reached (post-hoc LSD pairwise contrast, p < 0.05 for comparisons of days 1-2 to 7-8, p > 0.05 for all later time points). The analysis of interaction between age and social condition showed that differences between social conditions began on day 5 with the highest total secretion in QR workers and the lowest amount of secretion in QLBL workers (post-hoc LSD pairwise contrast, p < 0.05 for QR vs. QLBL, p > 0.05 for other comparisons, on days 11-12 p < 0.05 for all comparisons).

Overall, both ester and hydrocarbon amount increased with age but only the hydrocarbon amount differed significantly between the social conditions and covaried with oocyte size. Hydrocarbon amounts were highest in QR workers starting on days 5-6, in a similar fashion as the total secretion, and ester proportions covary with oocyte size but fluctuate over age groups and social conditions. To further examine the link between DG chemistry and the reproductive state of workers, we performed a discriminant analysis with three categorial classifications of the ovaries (see Methods for how categories were determined; Fig. 3). This analysis demonstrates that either esters, alkenes or the entire secretion provides more group differentiation than alkanes when workers are grouped according to ovarian activation.

### Examining the behavioral responses of *B. impatiens* workers to Dufour’s gland extracts

Due to the large impact of age we performed these experiments using same-aged workers at the age of 6-7 days from the two most distinct social conditions (QR and QLBL). Workers of this age showed significant differences in their ovary activation, amount and percent of esters and hydrocarbons (ovaries: Wilcoxon test, W = 4, p < 0.001; QR – 0.7 ± 0.04, QLBL – 1.9 ± 0.31; mean mm ± SE, n = 10), amount of esters (W = 0.8, p < 0.001; QR – 4.8 ± 1.1, QLBL – 0.9 ± 0.1; mean µg ± SE, n = 10), amount of hydrocarbons (W = 86, p = 0.005; QR – 14.4 ± 2.6, QLBL – 5.6 ± 1.1; mean µg ± SE, n = 10, percent esters (W = 79, p = 0.029; QR – 25.0 ± 2.2, QLBL – 19.3 ± 3.1; mean percent ± SE, n = 10; Fig. 4). In olfactometer bioassays, workers from both social conditions (QR, QLBL), given a choice between a callow worker DG extract (which served as a control) and QLBL DG extract, avoided QLBL DG extract (QLBL responders, n = 14, p = 0.01; QR responders n = 16, p = 0.004, binomial exact test; Fig. 5). When provided with a choice between callow extract vs. QR DG extract, workers of neither social conditions showed a significant preference (QLBL responder, n = 16, p = 1; QR responder, n = 16, p = 0.21). Similarly, when provided with a choice between QLBL and QR DG extracts, neither showed a significant preference (QLBL responder, n = 15, p = 1). Both same-age QR and QLBL workers preferred extracts that represented lack of competition (i.e., callow or queenright workers). On average, QLBL workers at the age of 6-7 days had lesser amounts of esters and higher ovary activation (Fig. 4), and preferred the control over QLBL extract and QLBL over QR extract, while same-age QR workers had, on average, more esters and hydrocarbons and lower ovary activation (Fig. 4), and preferred the control over either QLBL or QR extracts.

**Figure 5.**
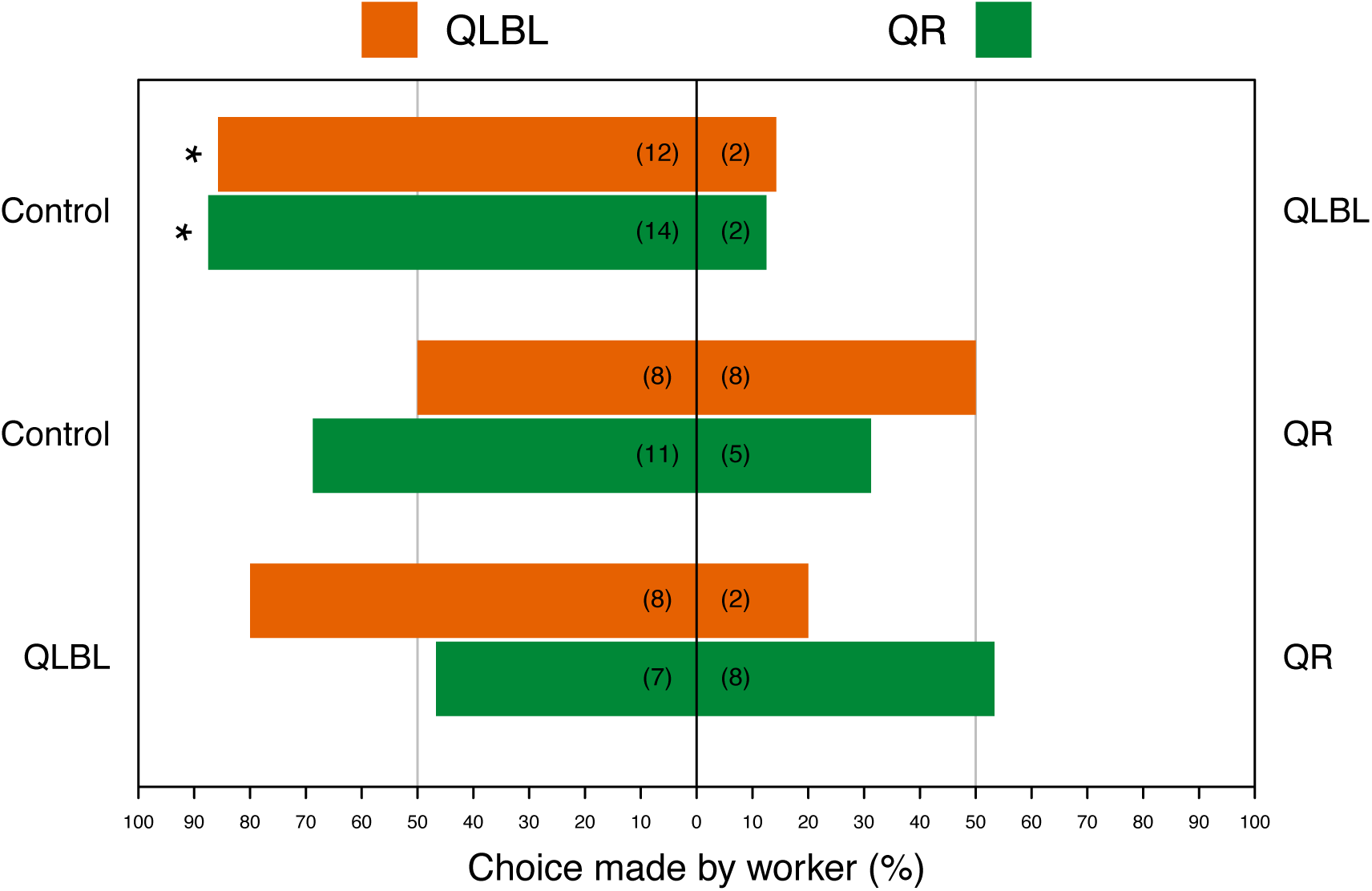
The proportion of queenless brood-less (QLBL) and queenright workers (QR) responding to 3 different combinations of stimuli, designated on the left and right of the plot: control vs. QLBL DG extract, control vs. QR DG extract, and QLBL vs. QR DG extract. Control stimuli consisted of callow worker DG extract. All stimuli were presented in the amount of 1 gland equivalent. The number of workers responding to each treatment is given in parenthesis on each bar. Orange bars represent QLBL response bees, and green bars represent QR response bees. Asterisks indicate significance (p <0.05) using exact binomial tests.

### Examining the antennal responses of *B. impatiens* workers to Dufour’s gland extracts

Electroantennogram recordings from antennae of 7-day-old workers of different social conditions responding to QLBL or QR DG extracts (also from 7-day-old workers) revealed that antennae of QLBL workers produced stronger responses, on average, than those of QR workers (p = 0.02, Fig. 6). Within social condition, QR workers produced stronger responses to DG extract from their own social condition than the QLBL extract (p < 0.001, Fig. 6), while QLBL workers showed no difference in response based on the extract type (p = 0.64). Overall, QLBL that had, on average, greater ovary activation and lesser amounts of esters (Fig. 4) exhibited stronger responses than QR, and QR that had, on average, lower ovary activation and more esters were more responsive to QR extracts.

**Figure 6.**
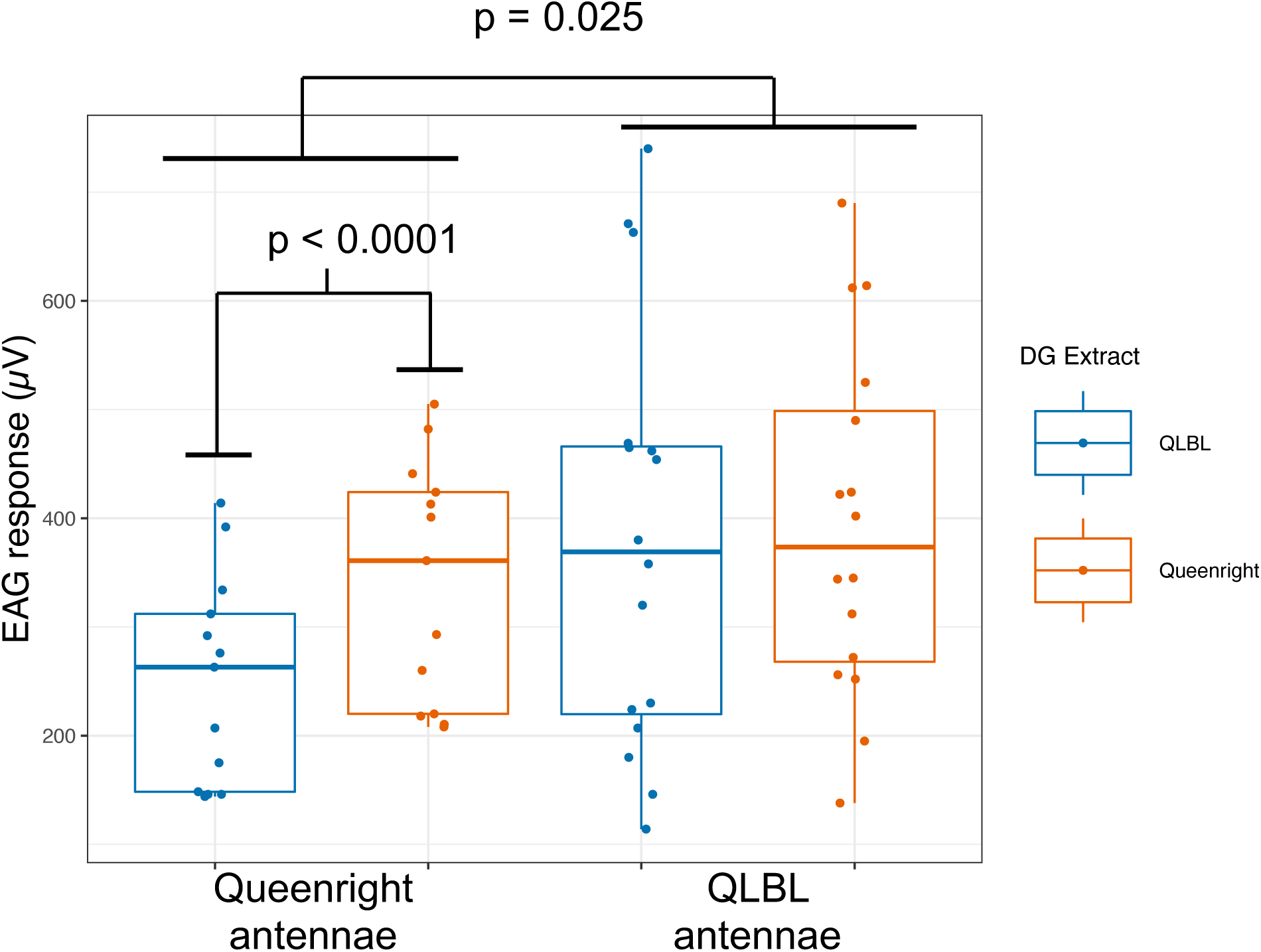
Electroantennogram responses (µV) of queenless brood-less and queenright worker antennae to Dufour’s gland extracts (1 worker equivalent) from queenless brood-less and queenright workers. Blue and orange boxplots represent responses to queenless brood-less and queenright Dufour’s gland extracts, respectively. Asterisks indicate significance (* p < 0.05, *** p < 0.001).

## Discussion

The results of this study reveal the presence of caste-specific and physiology-specific compounds in the DG of *B. impatiens* females. Our results further demonstrate that the DG secretion not only contains important information about caste, age, social, and reproductive condition, but also that the signaling properties of the gland contents are perceived and used by workers to discriminate among workers. This discrimination suggests that workers are attracted to DG extracts of workers that represent a lack of competition, and that QLBL workers, as opposed to QR workers, have higher sensitivity to DG extracts, representing an increased investment in sensory capability.The discriminant analysis of DG secretion showed clear differentiation of worker and queen groups, highlighting the caste-specific chemistry of the gland. This separation was primarily explained by worker-specific esters, gyne-specific esters, and queen-specific diterpene compounds, although alkanes and, to some extent, alkenes contributed to some of the variability (Fig. 1). The long chain esters that separate gynes from workers are also found in much lower amounts in queens, pointing towards a signal that functions either in caste recognition by workers, or sexual attraction of males. While the latter was not investigated in this study, caste recognition by workers is supported by characteristic behaviors of workers towards gynes. For example, although workers can be quite aggressive towards other workers or queens from a different colony, they remain docile in the presence of gynes (Pers. obs., ND and EA).

The diterpene compounds that characterize the queen DG secretion are primarily known from the labial glands of bumble bee males^44,45^, where they are hypothesized to serve as sex-pheromone components^46^. β-springene in particular has also been identified from the DG of a braconid wasp^47^, but the function it serves is unknown. Female sex pheromones in the related species *B. terrestris* were identified from head and body surface extracts of gynes^48^, but DG components have not been investigated as sex pheromones. Previous research comparing the DG secretion in unmated gynes of 5 species of European bumble bee found large variation in long-chain wax esters, with *B. pascuorum* and *B. lucorum* possessing up to 5% of hexadecenyl hexadecenoate or octadecyl octadec-9-enoate, while *B. hypnorum, B. lapidarius*, and *B. terrestris* had none or less 1%^49^. Interestingly, diterpenes were not found in the DG of these species, indicating that they may be species- and caste-specific. The DG’s diterpene compounds in *B. impatiens* could also function in caste recognition, a role that is often ascribed to cuticular hydrocarbons^50,51^. The DG of the honey bee contains ∼12 times more secretion in queens than workers and contains caste specific compounds, leading previous researchers to hypothesize that the gland deposits a signal of egg-maternity that distinguishes queen-from worker-laid eggs^52,53^. Caste-specific compounds, likely from the DG, have also been identified from the eggs of the bumble bee *B. terrestris*^54^. In *B. impatiens*, it is unclear why both the queen and worker castes would each need a unique set of compounds to differentiate egg-maternity, or why workers would advertise egg maternity, because this is likely to result in oophagy^3^, suggesting that these compounds might have another as yet unknown signaling function.

Workers produced a series of dodecyl esters that were not detected in extracts from gynes or mated queens (Fig S2, Table S2). These esters as well the hydrocarbon compounds in their DG may be used as signals by conspecific queens, gynes, males or workers. While we only focused on the signaling role of the secretion to other workers, the gland may contain multiple signals targeting different individuals, as occurs with CHCs which signal both fertility and nestmate identity in many social insects^51,55^, or with the honey bee queen mandibular pheromone, which functions as a primer pheromone to both workers and males, and a sex pheromone to males^56,57^. In *Bombus terrestris*, the main role of the DG worker-specific signals is in the interactions among workers^8,21,34,58^. QR workers may gain from both producing a signal that conveys harmony and respond to such a signal in order to maintain the social phase they are in (i.e., protect the queen and ensure the production of future gynes), while QL workers, who enter a reproductive mode, may gain from being tuned to signals that convey potential competition that may threaten the fate of their eggs. By manipulating social condition to produce either QLBL reproductive workers or QR non-reproductive workers, and then testing their perception and behavioral response to DG signals from those same groups, we show that workers preferentially avoid bees of a non-harmonious reproductive state (Fig. 5), with both QR and QLBL workers choosing callow over QLBL extract, and being indifferent between control and QR extracts. As bees from both social conditions were responding to aliquots of the same gland extract pooled from several colonies, nestmate information was obscured and is unlikely to explain the differences in attraction.

In electroantennography experiments, antennae from QLBL workers produced stronger responses than QR antennae (Fig. 6), suggesting that QLBL antennae are more sensitive to perceiving DG signals. Queenlessness is known to affect olfactory development and performance in the honey bee, where the worker olfactory system develops substantially during the first 6 days of life, and is strongly affected by the queen’s presence^59,60^. Bumble bees and honey bees have similar life span and 6-7 days is the same time period that workers in this study were kept queenless. Additionally, in honey bee workers, queenlessness is associated with a reduction in glomerular volume in the antennal lobe, and decreased learning of conditioned stimuli^61^. It is not known if similar processes govern bumble bee worker development. Reproductively active 7-day-old workers are likely capable of laying eggs and would benefit from detecting and avoiding the queen, who would aggress the workers that challenge her reproductive monopoly. Additionally, QR workers showed a reduced antennal response to QLBL extract in comparison to QLBL workers in electroantennography experiments (Fig 6). This could indicate that QR bees are operating in “harmony mode” within the colony with a higher threshold for detecting and aggressing other individuals. It is not known at what level this differential response is regulated. QR and QLBL bees could have differing numbers of olfactory receptors tuned to the DG compounds, or these chemical signals could be interpreted at higher levels of signal integration and processing in the brain.

Within workers, the composition of the DG secretion varied between social conditions, and the amount of the secretion increased with age (Fig. 2), with differences in the amounts and proportions of esters and hydrocarbons at certain age groups (eg, in 6-7-day-old QR and QLBL workers; Fig. 4). While the broad qualitative composition of DG extracts in *B. impatiens* workers is similar to related species of *Bombus*^8,54,62,63^, the influence of social condition and reproductive state on gland chemistry is different than the most closely studied species, *B. terrestris*. In *B. terrestris* workers, the percentages of DG esters are negatively correlated with ovarian activation^8,34^, and the size of the gland is positively correlated with ovarian activation^64^. Here, we demonstrate that in *B. impatiens*, the amounts of compounds of all chemical classes, including esters, are positively correlated with age but affected by social condition at specific ages, while the percent composition is more stable over time (Fig. 2B). This is distinctly different than the case of esters in *B. terrestris* and indicates that differences in the amounts of compounds observed at specific ages result from upregulation of the gland as a whole, instead of specific components changing independently. In some systems, when a worker transitions into fertility, their pheromonal secretion becomes more queenlike^33^. In *B. impatiens*, fertile worker secretions do not mimic queen secretions, instead both maintain the production of compounds unique to their own caste. Thus, the quantitative differences in DG components appears to represent a fertility signal unique to the worker caste as compared to *B. terrestris*, where the wax esters are only present in workers with inactivated ovaries and their absence fully mimics the queen secretion^8^. This may require a more finely tuned threshold for aggression and help explain the relatively docile conflict between queen and workers over reproduction in *B. impatiens* compared with *B. terrestris*^65,66^. Similar processes may take place during the competition phase when a large portion of the workers activate their ovaries. Furthermore, the DG esters also vary between the two species, with dodecyl esters predominating in *B. impatiens* and octyl esters predominating in *B. terrestris*^8^. Interestingly, dodecyl esters are also found in the labial glands of *B. terrestris* females^67^ whereas predominately octyl esters are found in the labial glands of *B. impatiens* (Orlova M, Villar G, Hefetz, A, Millar J, Amsalem E, in preparation), presenting an interesting case of inversion and suggesting a regulatory system that is shared but not identical in different bumble bee species.

Thus, our data suggest that the DG contains multiple concurrent signals that contain information about caste, age, social and reproductive condition. The presence of diterpene and ester compounds, which are much less common than aliphatic alkanes and alkenes in insects, may serve as additional channels with which to communicate this information. It is also possible that the signal carries different information to different perceiving individuals. For instance, diterpene compounds may signal “gyne” to conspecific workers and “mate” to conspecific males. Future research should investigate where compounds from the DG can be found in a colony, whether they are deposited onto eggs and/or the cuticular surface of the bee, and their role in worker-worker, worker-queen and queen-male interactions.

## Acknowledgments

The authors are grateful to Dr. Tom Baker for use of his electrophysiology equipment, and to Freddy Purnell for assistance in bumble bee colony maintenance and rearing.

## Author Contributions

EA, AH, GV designed the experiments. GV and ND performed dissections, ovary measurements, and gland extracts. ND and MO performed data and statistical analyses. GV performed olfactometer and GC-EAD bioassays. ND and JM performed GC-MS analysis. ND and EA wrote the manuscript. All authors provided feedback and edits.

## References

1. Orlova, M., Starkey, J. & Amsalem, E. A small family business: synergistic and additive effects of the queen and the brood on worker reproduction in a primitively eusocial bee. J. Exp. Biol. 223, (2020).

2. Starkey, J., Brown, A. & Amsalem, E. The road to sociality: brood regulation of worker reproduction in the simple eusocial bee Bombus impatiens. Anim. Behav. 154, 57–65 (2019).

3. Zanette, L. R. S. et al. Reproductive conflict in bumblebees and the evolution of worker policing. Evolution (N. Y). 66, 3765–3777 (2012).

4. Wenseleers, T. et al. A test of worker policing theory in an advanced eusocial wasp, Vespula rufa. Evolution 59, 1306–14 (2005).

5. Duchateau, M. J. & Velthuis, H. H. W. Ovarian development and egg-laying in workers of Bombus-terrestris. Entomol. Exp. Appl. 51, 199–213 (1989).

6. Hoover, S. E. R., Keeling, C. I., Winston, M. L. & Slessor, K. N. The effect of queen pheromones on worker honey bee ovary development. Naturwissenschaften 90, 477–80 (2003).

7. Vargo, E. L. & Laurel, M. Studies on the mode of action of a queen primer pheromone of the fire ant Solenopsis invicta. J. Insect Physiol. 40, 601–610 (1994).

8. Amsalem, E., Twele, R., Francke, W. & Hefetz, A. Reproductive competition in the bumble-bee Bombus terrestris: do workers advertise sterility? Proc. R. Soc. B 276, 1295–304 (2009).

9. Hefetz, A. The critical role of primer pheromones in maintaining insect sociality. Z. Naturforsch. C. 1–11 (2019). doi:10.1515/znc-2018-0224

10. Le Conte, Y. & Hefetz, A. Primer pheromones in social Hymenoptera. Annu. Rev. Entomol. 53, 523–542 (2008).

11. Oi, C. A. et al. The origin and evolution of social insect queen pheromones: Novel hypotheses and outstanding problems. BioEssays n/a-n/a (2015). doi:10.1002/bies.201400180

12. Kocher, S. D. & Grozinger, C. M. Cooperation, conflict, and the evolution of queen pheromones. J. Chem. Ecol. 37, 1263–1275 (2011).

13. Oi, C. A. et al. Dual effect of wasp queen pheromone in regulating insect sociality. Curr. Biol. 25, 1638–1640 (2015).

14. Smith, A. a., Hölldobler, B. & Liebig, J. Hydrocarbon signals explain the pattern of worker and egg policing in the ant Aphaenogaster cockerelli. J. Chem. Ecol. 34, 1275–1282 (2008).

15. Endler, A. et al. Surface hydrocarbons of queen eggs regulate worker reproduction in a social insect. Proc. Natl. Acad. Sci. U. S. A. 101, 2945–50 (2004).

16. Miller, D. G. & Ratnieks, F. L. W. The timing of worker reproduction and breakdown of policing behaviour in queenless honey bee (Apis mellifera L.) societies. Insectes Soc. 48, 178–184 (2001).

17. Dietemann, V., Peeters, C., Liebig, J., Thivet, V. & Hölldobler, B. Cuticular hydrocarbons mediate discrimination of reproductives and nonreproductives in the ant Myrmecia gulosa. Proc. Natl. Acad. Sci. U. S. A. 100, 10341–6 (2003).

18. Cuvillier-Hot, V., Cobb, M., Malosse, C. & Peeters, C. Sex, age and ovarian activity affect cuticular hydrocarbons in Diacamma ceylonense, a queenless ant. J. Insect Physiol. 47, 485–493 (2001).

19. Visscher, P. K. & Dukas, R. Honey bees recognize development of nestmates’ ovaries. Anim. Behav. 49, 542–544 (1995).

20. Smith, A., Hölldober, B., Liebig, J., Holldobler, B. & Liebig, J. Cuticular hydrocarbons reliably identify cheaters and allow enforcment of altruism in a social insect. Curr. Biol. 19, 78–81 (2009).

21. Amsalem, E. & Hefetz, A. The appeasement effect of sterility signaling in dominance contests among Bombus terrestris workers. Behav. Ecol. Sociobiol. 64, 1685–1694 (2010).

22. Fletcher, D. J. C. & Ross, K. G. Regulation of reproduction in eusocial Hymenoptera. Annu. Rev. Entomol. 30, 319–343 (1985).

23. Keeling, C. I., Plettner, E. & Slessor, K. N. Hymenopteran semiochemicals. Top. Curr. Chem. 239, 133–77 (2004).

24. Amsalem, E. One problem, many solutions: Female reproduction is regulated by chemically diverse pheromones across insects. in Advances in Insect Physiology (ed. Jurenka, R.) 59, 51 (Elsevier Ltd, 2020).

25. Mitra, A. Function of the Dufour’s gland in solitary and social Hymenoptera. J. Hymenopt. Res. 35, 33–58 (2013).

26. Abdalla, F. C. & Cruz-Landim, C. Dufour glands in the hymenopterans (Apidae, Formicidae, Vespidae): a review. Brazilian J. Biol. 61, 95–106 (2001).

27. Hefetz, A., M. Fales, H. & Batra, S. Natural polyesters: Dufour’s gland macrocyclic lactones form brood cell laminesters in Colletes bees. Science (80-.). 204, 415–417 (1979).

28. Cane, J. H. Chemical evolution and chemosystematics of the dufour’s gland secretions of the lactone-producing bees (Hymenoptera: Colletidae, Halictidae, and Oxaeidae). Evolution (N. Y). 37, 657–674 (1983).

29. Weseloh, R. M. Dufour’s gland: source of sex pheromone in a hymenopterous parasitoid. Science (80-.). 193, 695–697 (1976).

30. Pitts-Singer, T. L. et al. Comparison of the chemical compositions of the cuticle and Dufour’s gland of two solitary bee species from laboratory and field conditions. J. Chem. Ecol. 43, 451–468 (2017).

31. Shimron, O., Hefetz, A. & Tengo, J. Structural and communicative functions of Dufour’s gland secretion in Eucera palestinae (Hymenoptera; Anthophoridae). Insect Biochem. 15, 635–638 (1985).

32. Hefetz, A. Exocrine glands and their products in non-Apis bees: chemical, functional, and evolutionary perspectives. in Pheromone Communication in Social Insects (eds. Vander Meer, R. K., Breed, M. D., Espelie, K. E. & Winston, M. L.) (Westview Press, 1998).

33. Dor, R., Katzav-Gozansky, T. & Hefetz, A. Dufour’s gland pheromone as a reliable fertility signal among honeybee (Apis mellifera) workers. Behav. Ecol. Sociobiol. 58, 270–276 (2005).

34. Amsalem, E., Shpigler, H., Bloch, G. & Hefetz, A. Dufour’s gland secretion, sterility and foraging behavior: Correlated behavior traits in bumblebee workers. J. Insect Physiol. 59, 1250–1255 (2013).

35. Duchateau, M. J. & Velthuis, H. H. W. Development and reproductive strategies in Bombus terrestris colonies. Behavior 107, 186–207 (1988).

36. Amsalem, E., Grozinger, C. M., Padilla, M. & Hefetz, A. The Physiological and Genomic Bases of Bumble Bee Social Behaviour. Advances in Insect Physiology 48, (Elsevier Ltd., 2015).

37. Abdalla, F. C. & Cruz-Landim, C. da. Size differences in the Dufour gland of Apis mellifera Linnaeus (Hymenoptera, Apidae) between and within the female castes. Rev. Bras. Zool. 18, 119–123 (2001).

38. Starkey, J., Derstine, N. & Amsalem, E. Do bumble bees produce brood pheromones? J. Chem. Ecol. 45, 725–734 (2019).

39. Villar, G., Baker, T. C., Patch, H. M. & Grozinger, C. M. Neurophysiological mechanisms underlying sex- and maturation-related variation in pheromone responses in honey bees (Apis mellifera). J. Comp. Physiol. A 201, 731–739 (2015).

40. Park, K. C., Ochieng, S. a, Zhu, J. & Baker, T. C. Odor discrimination using insect electroantennogram responses from an insect antennal array. Chem. Senses 27, 343–52 (2002).

41. Aitchison, J. The Statistical Analysis of Compositional Data. (Chapman and Hall, 1986).

42. Martin, S. J., Takahashi, J.-I. & Katada, S.-I. Queen condition, mating frequency, queen loss, and levels of worker reproduction in the hornets Vespa affinis and V. simillima. Ecol. Entomol. 34, 43–49 (2009).

43. Derstine, N. T., Gries, R., Zhai, H., Jimenez, S. I. & Gries, G. Cuticular hydrocarbons determine sex, caste, and nest membership in each of four species of yellowjackets (Hymenoptera: Vespidae). Insectes Soc. 65, 581–591 (2018).

44. Bertsch, A., Schweer, H. & Titze, A. Chemistry of the cephalic labial gland secretions of male Bombus morrisoni and B. rufocinctus, two North American bumblebee males with perching behavior. J. Chem. Ecol. 34, 1268–1274 (2008).

45. Bertsch, A., Schweer, H. & Titze, A. Analysis of the labial gland secretions of the male bumblebee Bombus griseocollis (Hymenoptera: Apidae). Zeitschrift fur Naturforsch. - Sect. C J. Biosci. 59, 701–707 (2004).

46. Valterová, I., Martinet, B., Michez, D., Rasmont, P. & Brasero, N. Sexual attraction: A review of bumblebee male pheromones. Zeitschrift fur Naturforsch. - Sect. C J. Biosci. 74, 233–250 (2019).

47. Howard, R. W., Baker, J. E. & Morgan, E. D. Novel diterpenoids and hydrocarbons in the Dufour gland of the ectoparasitoid Habrobracon hebetor (Say) (Hymenoptera: Braconidae). Arch. Insect Biochem. Physiol. 54, 95–109 (2003).

48. Krieger, G. M. et al. Identification of queen sex pheromone components of the bumblebee Bombus terrestris. J. Chem. Ecol. 32, 453–71 (2006).

49. Chalikova, L., Hovorka, O., Ptacek, V. & Valterova, I. Exocrine gland secretions of virgin queens of five bumblebee species (Hymenoptera: Apidae, Bombini). Naturforsch. B 59c, 582–589 (2004).

50. Ren, W. et al. Maculatic Acids—Sex Attractant Pheromone Components of Bald-Faced Hornets. Angew. Chemie - Int. Ed. 57, 11618–11622 (2018).

51. van Zweden, J. S. & Ettorre, P. Nestmate recognition in social insects and the role of hydrocarbons. in Insect Hydrocarbons: Biology, Biochemistry, and Chemical Ecology (eds. Blomquist, G. J. & Bagnéres, A.-G.) 222–243 (Cambridge University Press, 2010).

52. Katzav-Gozansky, T., Soroker, V. & Hefetz, A. Plasticity of caste-specific Dufour’s gland secretion in the honey bee (Apis mellifera L.). Naturwissenschaften 84, 238–241 (1997).

53. Ratnieks, F. L. W. Egg-laying, egg-removal, and ovary development by workers in queenright honey bee colonies. Behav. Ecol. Sociobiol. 32, 191–198 (1993).

54. Ayasse, M. et al. Caste- and colony-specific chemical signals on eggs of the bumble bee, Bombus terrestris L. (Hymenoptera: Apidae). Chemoecology 9, 112–126 (1999).

55. Cournault, L. & de Biseau, J. C. Hierarchical perception of fertility signals and nestmate recognition cues in two dolichoderine ants. Behav. Ecol. Sociobiol. 63, 1635–1641 (2009).

56. Slessor, K. N., Winston, M. L. & Le Conte, Y. Pheromone communication in the honeybee (Apis mellifera L.). J. Chem. Ecol. 31, 2731–45 (2005).

57. Villar, G. & Grozinger, C. M. Primer effects of the honeybee, Apis mellifera, queen pheromone 9-ODA on drones. Anim. Behav. 127, 271–279 (2017).

58. Amsalem, E., Shamia, D. & Hefetz, A. Aggression or ovarian development as determinants of reproductive dominance in Bombus terrestris: interpretation using a simulation model. Insectes Soc. 60, 213–222 (2013).

59. Masson, C. & Arnold, G. Ontogeny, maturation and plasticity of the olfactory system in the workerbee. J. Insect Physiol. 30, 7–14 (1984).

60. Allan, S. A., Slessor, K. N., Winston, M. L. & King, G. G. S. The influence of age and task specialization on the production and perception of honey bee pheromones. J. Insect Physiol. 33, 917–922 (1987).

61. Morgan, S. M., Butz Huryn, V. M., Downes, S. R. & Mercer, A. R. The effects of queenlessness on the maturation of the honey bee olfactory system. Behav. Brain Res. 91, 115–126 (1998).

62. Tengö, J. A. N. et al. Species specificity and complexity of Dufour’s gland secretion of bumble bees. Comp. Biochem. Physiol. 99B, 641–646 (1991).

63. Hefetz, A., Taghizadeh, T. & Francke, W. The exocrinology of the queen bumble bee Bombus terrestris (Hymenoptera: Apidae, Bombini). Zeitschrift fur Naturforsch. Sect. C - J. Biosci. 51, 409–422 (1996).

64. Abdalla, F. C., Velthuis, H., Duchateau, M. J. & Cruz-Landim, C. D. Secretory cycle of the Dufour’s gland in workers of the bumble bee Bombus terrestris L. (Hymenoptera: Apidae, Bombini). Netherlands J. Zool. 49, 135–156 (1999).

65. Cnaani, J., Schmid-Hempel, R. & Schmidt, J. O. Colony development, larval development and worker reproduction in Bombus impatiens Cresson. Insectes Soc. 49, 164–170 (2002).

66. Padilla, M., Amsalem, E., Altman, N., Hefetz, A. & Grozinger, C. M. Chemical communication is not sufficient to explain reproductive inhibition in the bumblebee Bombus impatiens. R. Soc. Open Sci. 3, 160576 (2016).

67. Amsalem, E., Kiefer, J., Schulz, S. & Hefetz, A. The effect of caste and reproductive state on the chemistry of the cephalic labial glands secretion of Bombus terrestris. J. Chem. Ecol. 40, 900–912 (2014).

