## Supplementary Material for "Dufour’s gland analysis reveals caste and physiology specific signals in *Bombus impatiens*"

Supplementary Information

Figure S1 – Diagram of the two-choice olfactometer design. Olfactometers were fashioned from petri dishes (150 × 15 mm) where two equidistant holes (2 cm diameter) led to small plastic cups that held treatment or control stimuli on a glass slide. A 3 cm section of wide plastic straw was glued onto the bottom of the hole and extended into the cup, preventing re-entry into the main arena. Bees could not contact the glass slides unless they fully entered the cup, having fallen from the end of the plastic straw.

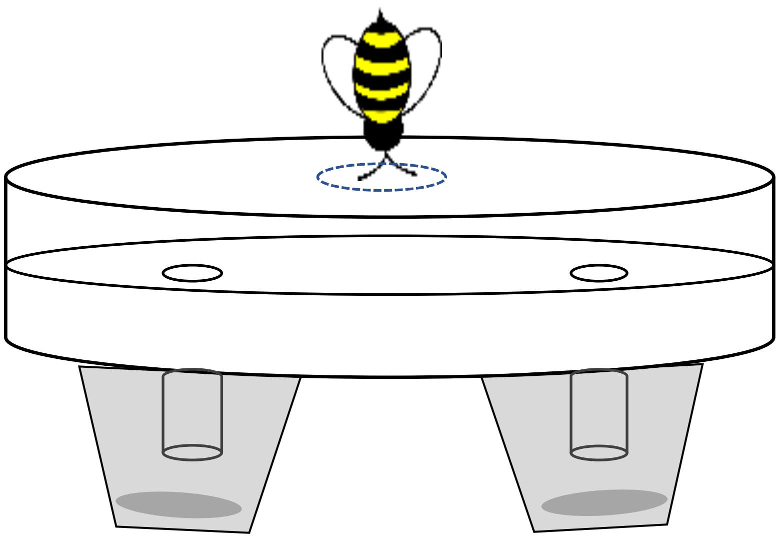

Figure S2 – Representative GC-FID chromatograms of *B. impatiens* workers, gynes and queens. Number labels correspond to compounds identified from hexane extracts of the Dufour's gland used in statistical comparisons. **1** – β-springene, **2** – springene isomer I, **3** – springene isomer II, **4** – eicosane IS ($C_{20}$), **5** – heneicosane ($C_{21}$), **6** – dodecyl-octanoate, **7** – docosane ($C_{22}$), **8** – tricosene ($C_{23:1}$), **9** – tricosane ($C_{23}$), **10** – dodecyl-decanoate, **11** – tetracosane ($C_{24}$), **12** – pentacosane ($C_{25:1}$), **13** - $C_{25}$, **14** – ester complex 1 (dodecyl-decanoate), **15** – heptacosene $C_{27:1}$, **16** – heptacosane $C_{27}$, **17** – ester complex 2 (hexadecyl decanoate), **18** – nonacosene I ($C_{29:1}$), **19** - nonacosene II ($C_{29:1}$), **20** – nonacosane ($C_{29}$), **21** – ester complex 3 dodecyl-9Z-hexadecenoate, **22** – hentriacontene I ($C_{31:1}$), **23** - hentriacontene II ($C_{31:1}$), **24** – hentriacontene ($C_{31}$), **25** - dodecyl-octadecenoate, **26** - ester complex 4 (octadecenyl tetradecanoate), **27** – ester complex 5 (octadecenyl hexadecanoate), **28** – terpene ester I, **29** – terpene ester II.

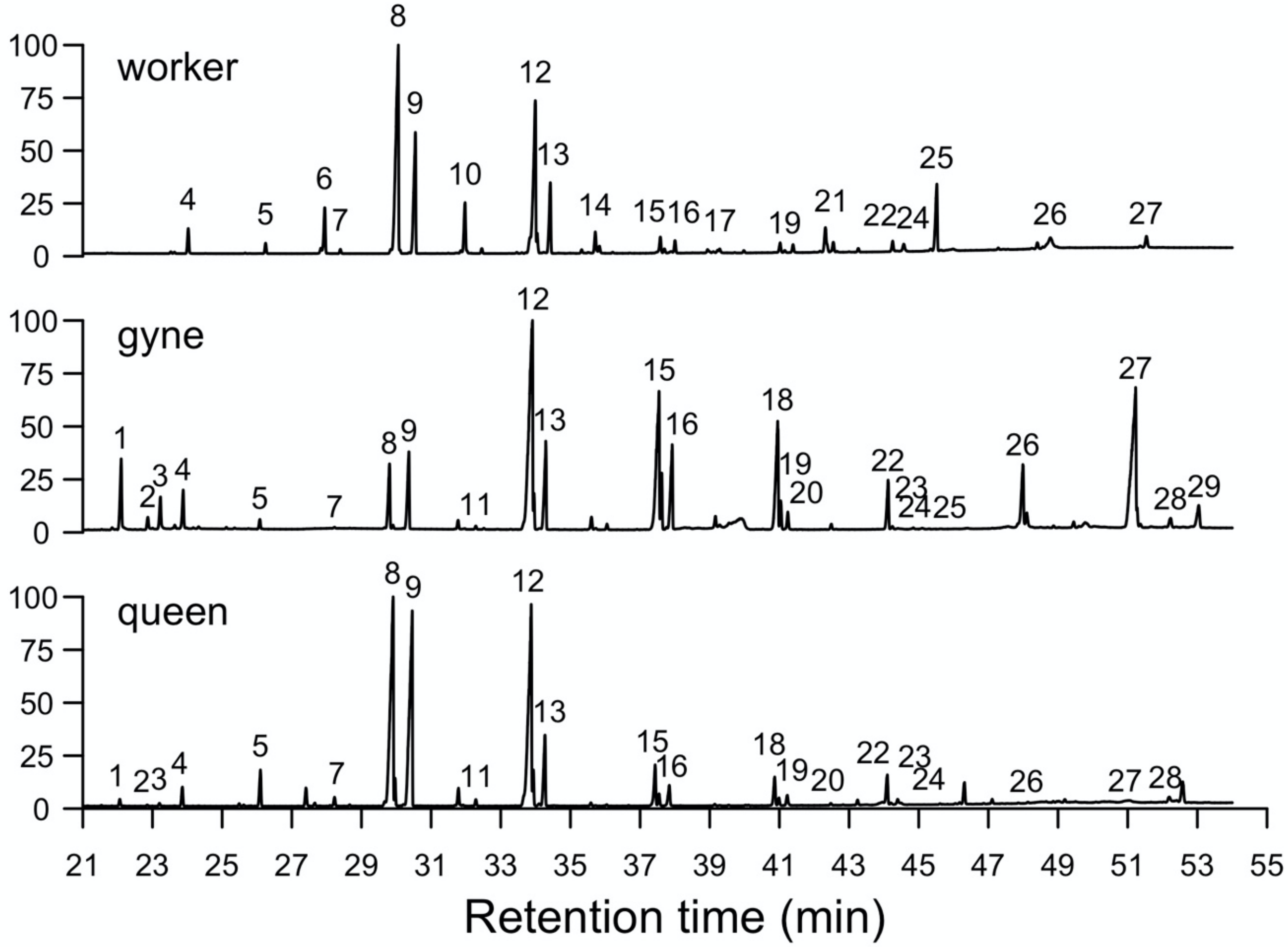

Figure S3 – Mean amount (µg) of the Dufour's gland chemical class "hydrocarbon" in 1-14 day old *B. impatiens* workers that were kept under three social condition treatments: queenless (QL), queenless and broodless (QLBL), or queenright (QR), shown as blue, orange, or green, respectively (n = 10 per treatment per day). The class hydrocarbon is separated into "alkane" and "alkene".

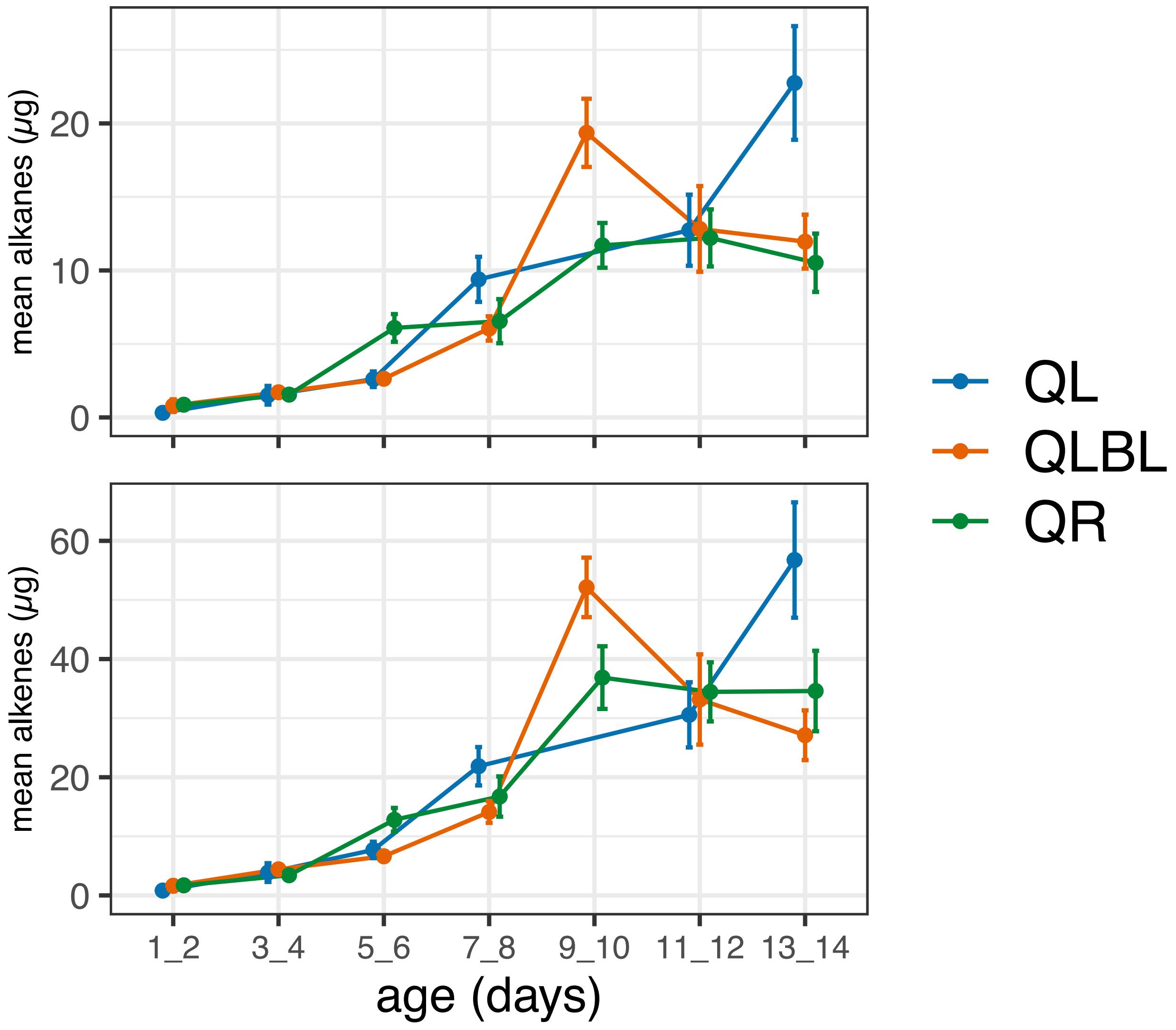

Table S1 - Summary of sample sizes per treatment and colony for *B. impatiens* workers, gynes, and queens used in chemical analyses of the Dufour's gland, olfactometer and EAG experiments in the current study. Colony was included in statistical models as a random factor.

| Experiment | Caste/treatment | Total sample size | Number of colonies |
|---|---|---|---|
| Chemical analysis of DG* | QL workers | 60 | |
| | QLBL workers | 70 | 7 |
| | QR workers | 69 | |
| | Gynes | 20 | 3 |
| | Queens | 20 | 20 |
| Olfactometer experiment** | QR workers | 47 | 11 |
| | QLBL workers | 40 | |
| | QR workers | 13 | 3 |
| EAG experiment | QLBL workers | 16 | |
| Total | | 315 workers<br>40 gynes/queens | 38 different colonies |

* The total sample size for workers does not include outliers that were excluded from the chemical analysis (n=11, see Results for an explanation)
** The total sample size does not include non-responsive workers (n=35, see Results for an explanation)

Table S2 – Relative abundances (percent of total secretion) and absolute amounts (µg) of Dufour's glands compounds used in discriminant analyses, along with Kovat's retention indices for mated queens (n = 20), virgin gynes (n = 20), and workers (n = 199) of all ages. Data are presented as means ± S.E.M

| Peak | Compound ID | Class | Retention Index (DB-5MS) | Queen relative amount | Gyne relative amount | Worker relative amount | Queen amount (µg) | Gyne amount (µg) | Worker amount (µg) |
|---|---|---|---|---|---|---|---|---|---|
| 1 | β-springene | diterpene | 1920 | 1.3 ± 0.33 | 3.2 ± 0.19 | 0 ± 0 | 1.82 ± 0.72 | 2.94 ± 0.4 | - |
| 2 | Springene isomer I | diterpene | 1953 | 0.2 ± 0.05 | 0.5 ± 0.03 | 0 ± 0 | 0.29 ± 0.11 | 0.43 ± 0.06 | - |
| 3 | Springene isomer II | diterpene | 1968 | 0.5 ± 0.14 | 1.3 ± 0.07 | 0 ± 0 | 0.71 ± 0.28 | 1.21 ± 0.16 | - |
| 4 | $C_{20}$ (internal standard) – Eicosane | alkane | 2000 | NA | NA | NA | NA | NA | NA |
| 5 | $C_{21}$ – Heneicosane | alkane | 2100 | 1.0 ± 0.11 | 0.7 ± 0.06 | 0.5 ± 0.01 | 1.51 ± 0.46 | 0.64 ± 0.08 | 0.18 ± 0.01 |
| 6 | Dodecyl-octanoate | ester | 2173 | - | - | 1.0 ± 0.07 | - | - | 0.36 ± 0.04 |
| 7 | $C_{22}$ – Docosane | alkane | 2200 | 0.3 ± 0.02 | 0.1 ± 0.01 | 0.2 ± 0.01 | 0.41 ± 0.11 | 0.1 ± 0.02 | 0.08 ± 0.01 |
| 8 | $C_{23:1}$ – Tricosene (two isomers) | alkene | 2275 | 23.1 ± 1.57 | 7.1 ± 0.52 | 22.5 ± 0.41 | 31.54 ± 8.53 | 6.46 ± 0.81 | 7.88 ± 0.57 |
| 9 | $C_{23}$ – Tricosane | alkane | 2300 | 15.6 ± 0.78 | 6.0 ± 0.26 | 10.6 ± 0.17 | 22.73 ± 6.37 | 5.19 ± 0.57 | 3.79 ± 0.28 |
| 10 | Dodecyl-decanoate | ester | 2369 | - | - | 2.6 ± 0.08 | - | - | 0.89 ± 0.07 |
| 11 | $C_{24}$ – Tetracosane | alkane | 2400 | 0.4 ± 0.02 | 0.2 ± 0.01 | 0.4 ± 0.01 | 0.55 ± 0.15 | 0.13 ± 0.01 | 0.14 ± 0.01 |
| 12 | $C_{25:1}$ – Pentacosene (two isomers) | alkene | 2477 | 31.8 ± 1.14 | 28.2 ± 0.84 | 27.7 ± 0.36 | 51.35 ± 15.34 | 24.43 ± 2.61 | 9.46 ± 0.7 |
| 13 | $C_{25}$ – Pentacosane | alkane | 2500 | 6.5 ± 0.41 | 4.2 ± 0.18 | 8.1 ± 0.15 | 10.42 ± 3.31 | 3.43 ± 0.29 | 2.59 ± 0.19 |
| 14 | Dodecyl dodecanoate (ester complex 1*) | ester | 2567 | - | - | 1.8 ± 0.08 | - | - | 0.66 ± 0.07 |
| 15 | $C_{27:1}$ – Heptacosene | alkene | 2673 | 5.0 ± 0.69 | 12.6 ± 0.3 | 3.4 ± 0.1 | 14.13 ± 6.61 | 10.41 ± 0.96 | 1.24 ± 0.12 |
| 16 | $C_{27}$ – Heptacosane | alkane | 2700 | 1.7 ± 0.25 | 3.5 ± 0.23 | 1.4 ± 0.04 | 4.71 ± 2.23 | 2.78 ± 0.23 | 0.5 ± 0.05 |
| 17 | Hexadecyl decanoate (ester complex 2*) | ester | 2769 | - | - | 1.5 ± 0.13 | - | - | 0.55 ± 0.06 |
| 18 | $C_{29:1}$ – Nonacosene I | alkene | 2874 | 2.7 ± 0.39 | 5.4 ± 0.53 | 1.2 ± 0.04 | 7.02 ± 3.47 | 3.98 ± 0.37 | 0.56 ± 0.06 |
| 19 | $C_{29:1}$ – Nonacosene II | alkene | 2881 | 1.0 ± 0.25 | 1.9 ± 0.4 | 0.5 ± 0.02 | 3.56 ± 2.04 | 1.9 ± 0.64 | 0.18 ± 0.02 |
| 20 | $C_{29}$ – Nonacosane | alkane | 2900 | 1.8 ± 0.31 | 0.9 ± 0.12 | 0.8 ± 0.02 | 2.05 ± 0.65 | 0.67 ± 0.09 | 0.28 ± 0.02 |
| 21 | Dodecyl hexadecenoate (ester complex 3*) | ester | 2951 | 0.1 ± 0.02 | - | 3.1 ± 0.16 | 0.03 ± 0.01 | 0 ± 0 | 1.37 ± 0.21 |

| | | | | | | | | | |
|---|---|---|---|---|---|---|---|---|---|
| 22 | $C_{31:1}$ – Hentriacontene I | alkene | 3075 | 3.0 ± 0.45 | 2.1 ± 0.13 | 1.2 ± 0.06 | 5.59 ± 3.12 | 1.67 ± 0.16 | 0.65 ± 0.08 |
| 23 | $C_{31:1}$ – Hentriacontene II | alkene | 3083 | 0.4 ± 0.24 | 0.1 ± 0.01 | 0.2 ± 0.02 | 1.92 ± 1.66 | 0.07 ± 0.01 | 0.12 ± 0.02 |
| 24 | $C_{31}$ – Hentriacontane | alkane | 3100 | 1.7 ± 0.44 | 0.03 ± 0.01 | 0.4 ± 0.02 | 0.77 ± 0.13 | 0.02 ± 0.01 | 0.15 ± 0.01 |
| 25 | Dodecyl octadecenoate | ester | 3155 | 0.04 ± 0.01 | 0.01 ± 0.01 | 5.1 ± 0.29 | 0.03 ± 0.01 | - | 1.59 ± 0.14 |
| 26 | Octadecenyl tetradecanoate (ester complex 4*) | ester | 3333 | 0.3 ± 0.1 | 5.4 ± 0.24 | 2.4 ± 0.16 | 0.53 ± 0.32 | 4.4 ± 0.37 | 1.16 ± 0.17 |
| 27 | Octadecenyl hexadecenoate (ester complex 5*) | ester | 3539 | 0.5 ± 0.25 | 13.2 ± 1.05 | 3.3 ± 0.27 | 1.7 ± 1.02 | 10.38 ± 0.97 | 1.12 ± 0.12 |
| 28 | Terpene ester I | ester | 3601 | 1.0 ± 0.21 | 1.2 ± 0.18 | - | 0.82 ± 0.31 | 1.17 ± 0.21 | - |
| 29 | Terpene ester II | ester | 3643 | 0.1 ± 0.04 | 2.2 ± 0.26 | - | 0.18 ± 0.13 | 2.13 ± 0.35 | - |

* Ester complexes consisted of multiple often overlapping or co-eluting esters that were treated as a single variable. The most prevalent compound in the complex is given as the name. Other compounds in a given complex are esters with the same total number of carbons as the predominate compound, but different chain lengths of the acid or alcohol portion.
